# Continuous Polony Gels for Tissue Mapping with High Resolution and RNA Capture Efficiency

**DOI:** 10.1101/2021.03.17.435795

**Authors:** Xiaonan Fu, Li Sun, Jane Y. Chen, Runze Dong, Yiing Lin, Richard D. Palmiter, Shin Lin, Liangcai Gu

## Abstract

Current technologies for acquiring spatial transcript information from tissue sections rely on either RNA probes or spatial barcodes. The former methods require a priori knowledge for probeset formulation; the latter have yet to achieve single cell resolution and/or transcript capture efficiencies approaching dissociative, single-cell methods. Here, we describe a novel spatial transcriptome assay called <u>p</u>olony (or DNA cluster)-<u>i</u>nde<u>xe</u>d <u>l</u>ibrary-sequencing (PIXEL-seq). It improves upon other spatial barcoding methods by employing “continuous” polony oligos arrayed across a customized gel surface. In terms of assay performance, PIXEL-seq attains ≤ 1 µm resolution and captures >1,000 unique molecular identifiers/10×10 µm^2^. In other words, this global, naive platform achieves subcellular spatial transcriptome mapping while maintaining high transcript capture efficiencies.

---

The spatial organization and dynamics of cell-specific gene expression in tissues are fundamental to the understanding of molecular aspects of health and disease. Spatially resolved transcriptome profiling has been achieved by imaging-based *in situ* RNA hybridization (*1-3*) and sequencing (*4, 5*) and oligonucleotide array-indexed (or spatially barcoded) RNA sequencing (*6-9*). Technologies for spatially barcoded RNA sequencing have employed arrayed oligos with spatially resolved indices (e.g., spotted (*6*), self-assembled bead (*7, 8*), and *in situ* synthesized (*9*) arrays) as primers. These oligos are incorporated into in situ synthesized complementary DNAs (cDNAs) derived from a tissue section placed atop the array. Subsequent sequencing of the indexed cDNAs reveals the RNA transcripts and their spatial locations. Compared with imaging methods, spatial barcoding methods are not restricted by the diffraction-limited sampling of highly crowded molecules in cells and do not require specialized equipment and expertise for end users. However, due to limitations of existing oligo arrays, the reported methods do not have single-cell resolution and/or high RNA capture efficiency achieved by tissue dissociative, single-cell RNA sequencing (or scRNA-seq) (*10, 11*).

To capture transcripts in a tissue section, an ideal oligo array should have small features, like pixels of a camera sensor, with minimal feature-to-feature gaps and a homogenous oligo distribution easily accessible to tissue transcripts (hereafter termed a “continuous” distribution). Traditional microarrays (*12*) were developed mainly for analyzing homogenized samples, for example, mixed nucleic acids in a solution, not morphologically intact tissues. To avoid DNA diffusion between features, they have been fabricated with discrete features (or spots) often showing inconsistent DNA densities across substrate surfaces, resulting in biases in spatial RNA capturing. In next-generation sequencing, ultra-dense arrays are formed with polymerase colonies (known as polonies (*13, 14*) or DNA clusters (*15*)) and rolling circle colonies (rolonies or DNA nanoballs (*16*)) with diameters of ∼1 µm or less. Polony or rolony DNA might be converted to capture oligos bearing defined 3’ priming sequences. These oligo arrays with features ≥ 10-fold smaller than mammalian cells (typically 10 to 100 µm in diameter) could provide a sufficient resolution for segmenting spatially indexed cDNAs into individual cells. However, polonies and rolonies produced in available Illumina and other sequencing flow cells are only customized for DNA sequencing, not spatial barcoding. Indeed, like traditional arrays, they all have discrete and non-homogeneous DNA distributions. The substrates used in the flow cells (e.g., glass) might not provide sufficient constraints on tissue RNA diffusion, an increasingly detrimental factor for the detection at a higher resolution.

Here, we developed <u>p</u>olony-inde<u>xe</u>d library-sequencing (PIXEL-seq) using polony-derived oligo gel arrays for the spatial barcoding (**Fig. 1a**). By screening polyacrylamide (PAA) gel fabrication conditions, we found that oligo arrays derived from polonies formed on a crosslinked PAA gel surface (hereafter named “polony gel”) are ideal for the tissue barcoding applications. We first confirmed that widely used polonies grown in porous substrates such as a semifluidic linear PAA gel in Illumina sequencing flow cells (*15*) show typical discrete features with the highest intensities at polony centers and the lowest at the borders (**Fig. 1b**, left panel). In next-generation sequencing, polony peaks facilitate the registration of fluorescence signals, but might cause inconsistency in RNA capturing. Interestingly, by modifying the gel casting condition, continuous polonies were formed on the crosslinked PAA surface (**Fig. 1b**, left panel). These polonies show relatively fast size expansion and a more homogenous DNA distribution than discrete ones, likely because the polony formation close to the gel surface has less constraint on the amplification reaction.Although continuous polonies are connected, they rarely interpenetrate, and their borders can be delineated by the sequencing (**Fig. 1b**, right panel). Notably, compared with discrete polonies grown inside of PAA gels, continuous polonies can be efficiently cleaved by restriction digestion; 93.6% of double-stranded polony DNAs were digested to expose a 3’ priming site for cDNA synthesis by the *Taq*I digestion (**fig. S1**). Given the above advantages, continuous polonies are better suited than the Illumina polonies for creating high-quality oligo gels for RNA capturing.

**Figure 1.**
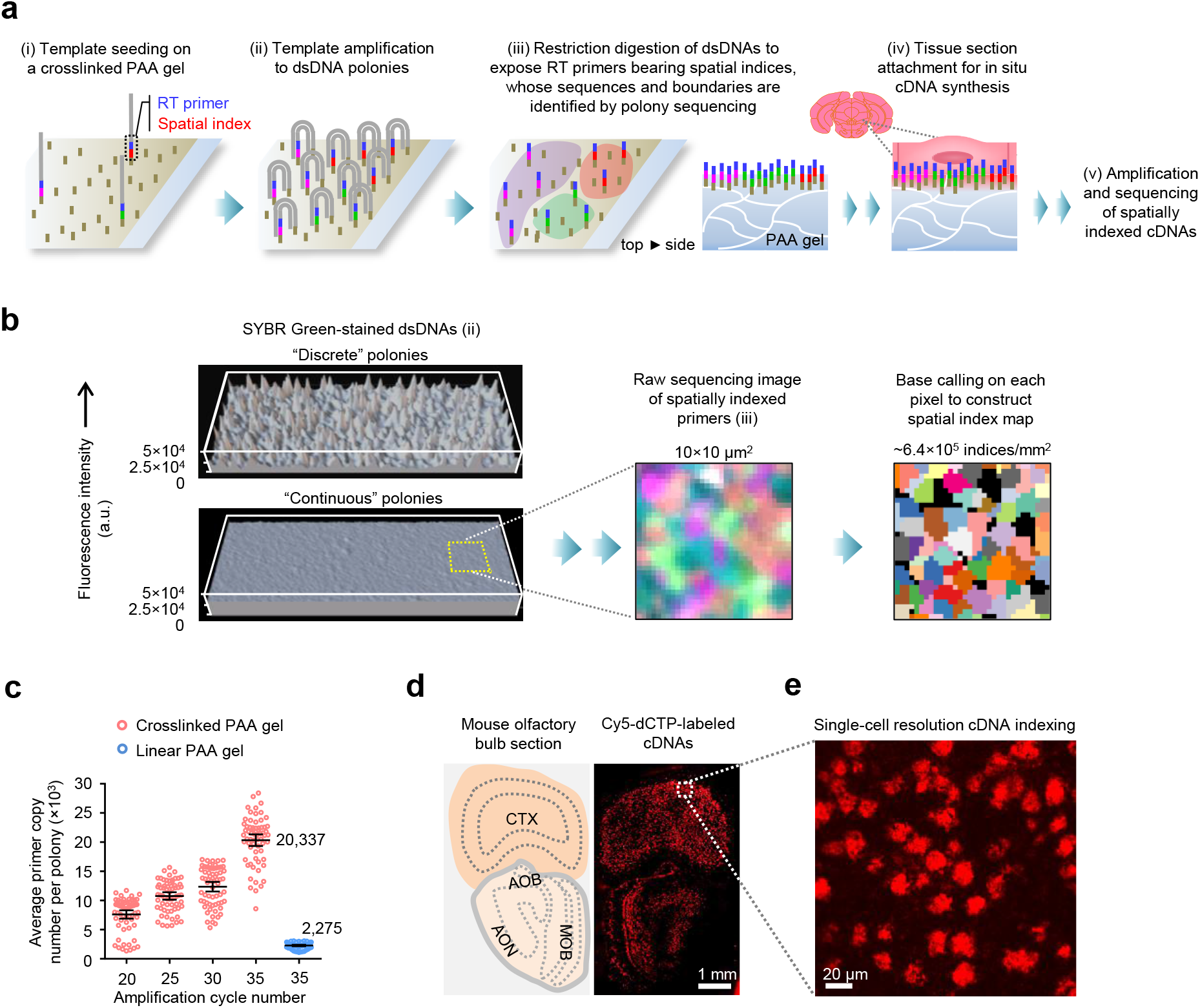
Principle of PIXEL-seq. (**a**) Schematic of PIXEL-seq-based spatial transcriptome analysis. 368-bp DNA templates bearing 24-bp random barcodes, a 20-bp poly(T) probe, and two *Taq*I restriction sites are amplified to polonies on a PAA gel surface. Barcode sequences and spatial coordinates were determined by polony sequencing. A tissue cryosection is placed onto a polony gel for mRNA capture and spatial indexing prior to library amplification for next-generation sequencing. (**b**) 3D intensity profiles of discrete and continuous polonies stained by SYBR Green. Polony sequencing images are converted to a spatial index map by pixel-level base calling. (**c**) Quantification of DNA copies per polony by real-time image quantification. (**d**) Cy5-labeling of spatially indexed cDNAs in a mouse olfactory bulb section. (**e**) Labelled cDNA signals show single-cell resolution.

To manufacture polony gels with spatially defined indices, we amplified ∼370-base pair (bp) templates bearing 24-bp spatial indices randomly seeded on a gel with a customizable size (e.g., 6 × 30 mm^2^) casted on a coverslip and then assembled in a flowcell, similarly to what we previously reported (*14*). Spatial indices were sequenced by a sequencing-by-synthesis chemistry (*15*) via an in-house constructed apparatus. Raw sequencing images were converted to a pixel-level spatial index map (**Fig. 1b**, right panel) by a base-calling pipeline which determines the major index species in each image pixel (0.325 × 0.325 µm^2^) (**fig. S2**).

To assess the resolution of our polony gels, we measured feature densities and sizes. We typically identified ∼0.5-0.8 million unique indices (or features) per mm^2^, close to the polony density of an Illumina non-patterned HiSeq flowcell. The average feature size is 1.17 ± 0.1 µm^2^ and can slightly increase at a lower feature density (**fig. S3a** and **3b**). Compared with other oligo arrays used by the reported methods (*6-8*), our feature sizes are ∼3 to 200-fold finer (**table S1**). The decreased feature size offers the sufficient resolution for image-guided cell segmentation for mammalian tissues; for example, our computational simulation suggests that, in an array covered by a tissue section with the average cell diameter of 15.9 ± 4 µm, the percentage of features contacting only one cell increases from 50.3 to 96.3% if the feature diameter decreases from 10 to 1 µm (**fig. S4**).

The RNA capture efficiency correlates to the density of array oligos and their accessibility to tissue RNAs. Crosslinked PAA gels have been reported to exhibit a 10 to 30-fold higher oligo capturing efficiency than solid surface substrates such as glass slides (*17*). Compared with linear PAA used in the Illumina sequencing method (*15*), the crosslinked gel provides greater mechanical strength and stability for tissue mapping assays. We demonstrated that it supports highly efficient polony amplification, yielding an average 20,337 template copies per feature after 35 amplification cycles (**Fig. 1c** and **fig. S5**), a ∼9-fold increase from that of a reported Illumina method (*13*). This process produces an oligo density of ∼1.74 × 10^4^ molecules/µm^2^, which is close to that of total oligos spotted on a glass substrate (*6*) (**table S1**). Our polony gels typically have over 85% substrate surface covered with DNAs (**fig. S3c**). A higher coverage can be achieved by increasing the template concentration and a higher resolution sequencing image setup.

After exposing a 3’ oligo(dT) sequence by the *Taq*I digestion, we tested the resolution and capturing efficiency of global messenger RNA (mRNA) detection. We first focused on cryosectioned tissues without fixation or other pretreatments, with the goals of avoiding adverse effects of fixatives on cDNA synthesis and having a simplified protocol. Our protocol was to attach 10-μm frozen sections to dried gels and then add reagents for cDNA synthesis. Fluorescence imaging of Cy5-labelled cDNAs in a mouse olfactory bulb-isocortex section achieved clear single-cell resolution (**Fig. 1d and 1e**), implying minimal lateral diffusion. We further assessed the impact of the diffusion by registering images of a nuclear stained olfactory bulb section on a dried gel as the reference and the synthesized cDNAs; a scaling factor was determined to be 1.028 ± 0.016 (**fig. S6**). Our results suggest that the PAA substrate can constrain the RNA diffusion to preserve the spatial resolution (**fig. S7a**). Additionally, the crosslinked PAA gave a 17.6-fold higher Cy5-labelled cDNA signal than the linear PAA (**fig. S8**), probably due to a higher oligo accessibility on the crosslinked gel surface (**fig. S7b**).

As a proof-of-principle for subcellular spatial transcriptome mapping, we analyzed mouse accessory olfactory bulb sections by PIXEL-seq. A polony gel with a sequenced feature density of ∼0.5 million per mm^2^ (**fig. S9a**) was attached to the tissue section, and the spatially indexed library was analyzed by next-generation sequencing. Guided by the predetermined spatial index map, sequencing reads were constructed into a digital spatial transcriptome image at subcellular resolution (**fig. S9b**). Each segmented cell was estimated having ∼50 clustered indices. The median unique molecular identifiers (UMIs) per bin (10 × 10 µm^2^) was ∼1,000, while DBSCAN segmented cell was assigned with more UMIs (median = 1,800) (**fig. S9c**).

To further assess the performance of PIXEL-seq, we applied the assay to a main olfactory bulb (MOB) section of the mouse brain, as was done by previous global, naive spatial technologies (*8, 18, 19*). MOB sections have relatively dense neuronal cell bodies from distinct neuron types presented in multiple morphological layers (*8*). We sought to determine whether our data can show expected layers and other histological features. We analyzed three replicate sections and tested the performance by two steps: i) unsupervised spatial pattern detection with high-resolution feature sizes, and ii) generating high-resolution spatial expression patterns of individual genes. Again, we confirmed the high resolution and sensitivity of mRNA detection (**Fig. 2a**). ∼23,000 unique genes are confidently detected in at least one replicate (reads count > 10; **fig. S10a**). 78.3% of all the genes were mapped to protein coding regions. A high correlation between each replicate indicates the robustness of PIXEL-seq (**Fig. S10b-c**). The unsupervised analysis by Seurat (*20*) identified ten distinct spatial varied clusters, in coordination with the known anatomical structures including olfactory nerve layers (ONLs), a glomerular layer (GL), an external plexiform layer (EPL), a granular cell layer externa (GCL-E), a granular cell zone deep (GCL-D), and a subependymal zone (SEZ) (**Fig. 2b**). Our result for layer-specific expression signatures agrees with their corresponding *in situ* hybridization (ISH) images from the Allen Brain Atlas (e.g. *Fabp7, Penk*, and *Doc2g*) (**Fig. 2c**). With a sequencing depth of 81.9 ± 7.6% on the MOB libraries, PIXEL-seq generates on average 58.9 UMIs at 2 µm resolution, 1,199 at 10 µm resolution, and 25,618 at 50 µm resolution (**Fig. 2d** and **fig. S11**). In comparison to existing technologies on the same tissue at 10 µm resolution, HDST captures 12 UMIs (*8*); Slide-seqV2, 494 UMIs (*18*). Upon analyzing a large bin size of 50 µm, PIXEL-seq captures 25,618 UMIs, which compares favorably to the 15,377 UMIs detected at 55-µm feature (not considering feature-to-feature gaps) by a recently updated Visium method. The Visium dataset on MOB was reanalyzed from a recent preprint (*21*).

**Figure 2.**
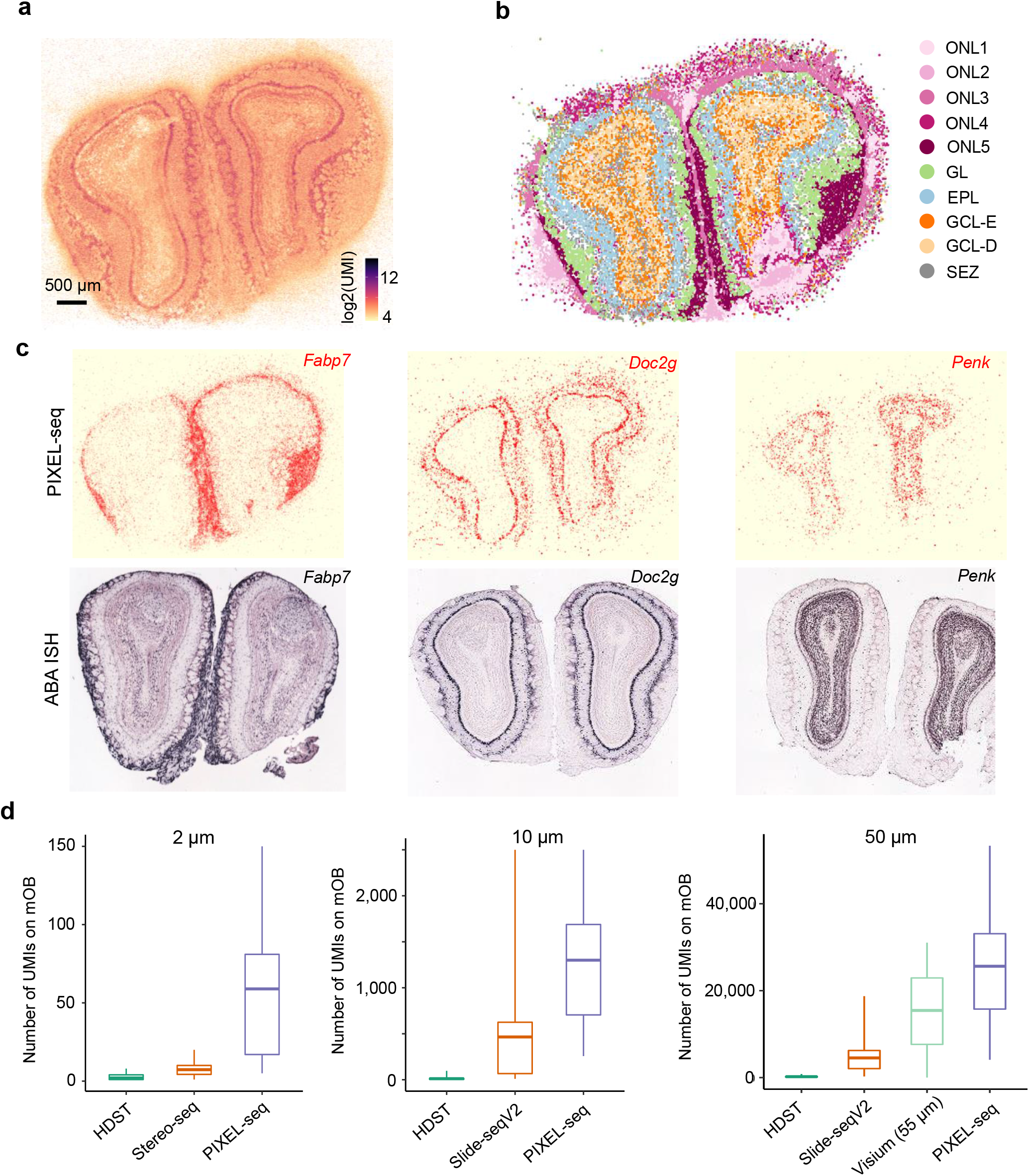
Spatial transcriptome profiling of the mouse main olfactory bulb (mOB). (**a**) Spatial distribution of gene density per polony (1 pixel = 0.325 × 0.325 μm^2^). (**b**) Unsupervised clustering by Seurat at binned feature size of 25 μm. (**c**) Examples of found genes showing spatial patterns consistent with the Allen ISH data. (d) Boxplots of mOB UMIs/bin captured by PIXEL-seq and other similar methods at the indicated resolution: 2 μm (grouping 7 pixels); 10 μm (32 pixels); 50 μm (154 pixels).

We next applied PIXEL-seq to analyze tissue samples which require a high mRNA capture efficiency at subcellular resolution. One region of interest is the parabrachial nucleus (PBN), which is located in the pons and relays sensory information from the periphery to the forebrain. Functional studies have shown that the PBN is important for responding to both internal and external stimuli, as well as maintaining homeostasis. All major sensory systems can activate neurons in the PBN, but specific neural circuits involved have not been well established. Recent studies have identified unique markers that label neurons involved in distinct behavior responses including *Penk* and *Pdyn* for thermal sensation (*22*), *Tac1* for pain (*23*), and *Calca* for appetite, visceral malaise, and threat detection. To identify unique PBN neuron markers that may distinguish responses to different sensory stimuli, we first investigated the overall spatially patterned genes through PIXEL-seq on three sections spanning the middle to caudal PBN from a wildtype adult mouse. An immediate overview of spatially detected gene profiles reveals abundant reads grouping into cell shape (**Fig. 3a**).While checking pattern of known PBN neuron markers, we found *Calca* and *Tac1* have a relative high expression and are spatially clustered whereas *Penk* is only partially overlapped with *Calca* and *Tac1* (**Fig. 3a**). These observations are confirmed by single molecule fluorescence *in situ* hybridization (smFISH), which shows the similar expression pattern and gene abundance level where *Tac1* and *Calca* are colocalized and largely do not overlap with *Pdyn* or *Penk* (**Fig. 3b**).

**Figure 3.**
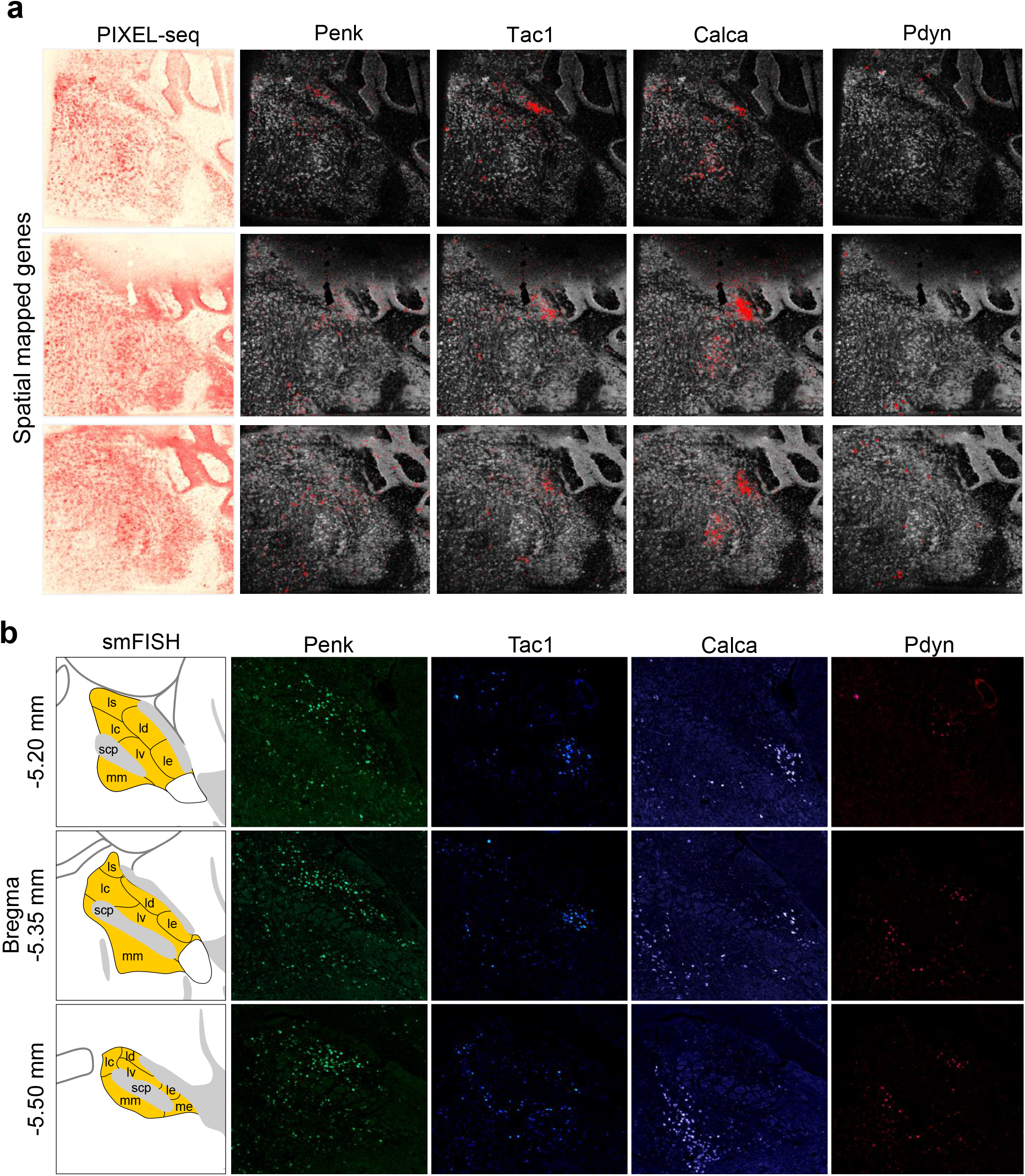
PIXEL-seq analysis of spatial patterns of selected PBN marker genes. (**a**) Spatial distribution of total UMI density and four marker genes (*Penk, Tac1, Calca*, and *Pdyn*) in anterior, middle, and posterior PBN sections. Spots in gray represent the overall UMI distribution and spots in red denote the maker gene read signals. (**b**) Validation of the PBN marker gene expression by hybridization chain reaction (HCR) FISH.

Particularly, the unsupervised spatial clustering by Seurat identified the PBN as the Cluster 6 (**Fig. 4a-c**). A subcellular-resolution mapped read image shows that some of the *Calca* reads are more enriched at the boundary of cells (**Fig. 4d-e**). These results provide a chance to explore the distinct regulatory mechanisms for PBN sensory functions.

**Figure 4.**
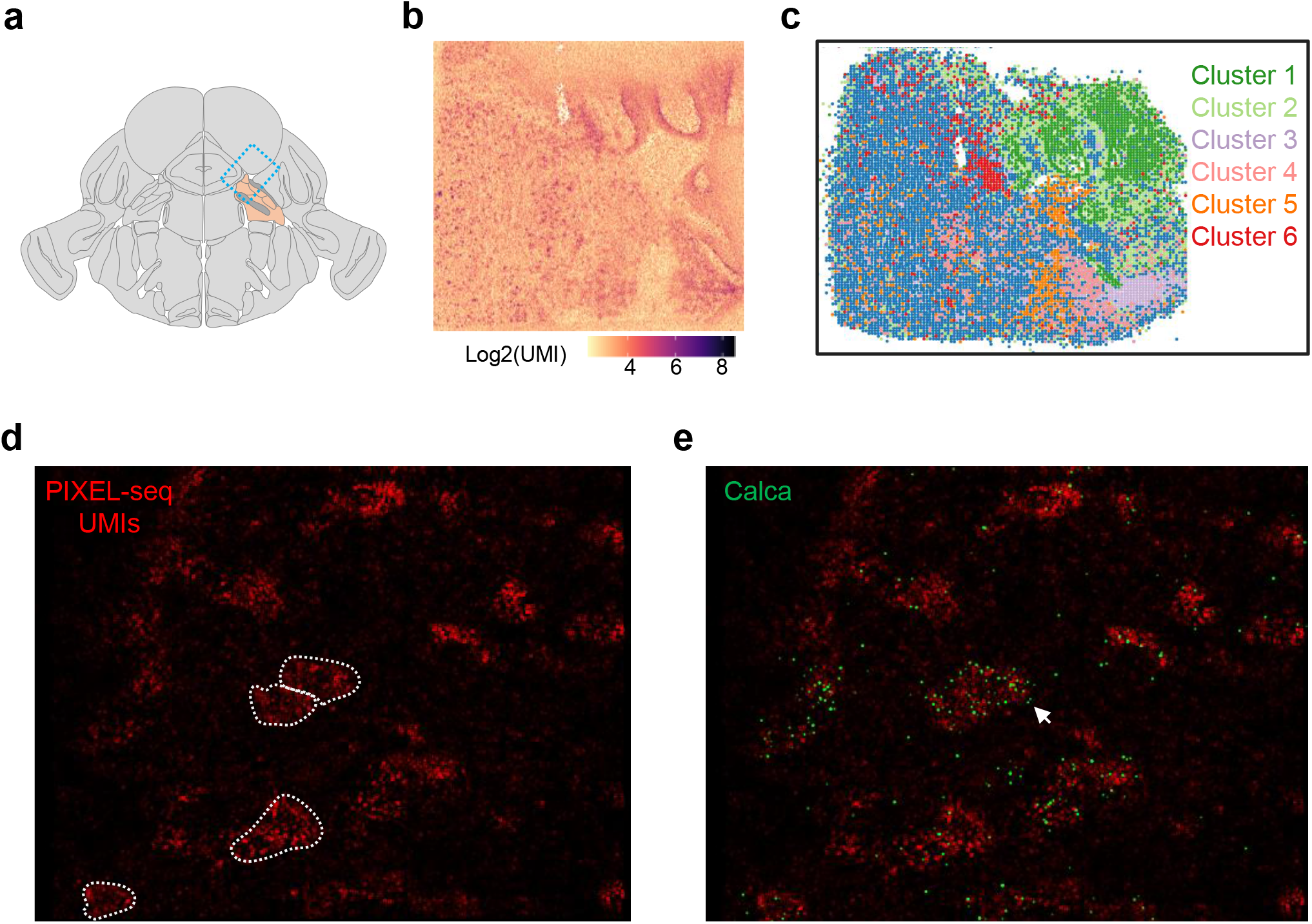
Spatial gene expression profile of PBN at subcellular resolution. (**a**) Illustration of the PBN cryosection site (in orange) and direction in the mouse brain. (**b**) Spatial distribution of gene density per feature (feature size = ∼1μm). (**c**) Unsupervised clustering of PIXEL-seq data on the PBN section by Seurat at a binned feature size of 25 μm. The Cluster 6 is overlapped with CGRP neurons (**d**) Gene abundance distribution at subcellular resolution. Selected cells showing *Calca* signals were manually segmented as references. (**e**) Subcellular distribution of *Calca* signals (in green).

Although PIXEL-seq was only tested for the poly(T) oligo-captured global transcriptome profiling, it should be applicable to other spatial barcoding-based “omics” assays. Restriction digested polony-oligos can be primed with any defined 3’ adapter sequences (e.g., random or gene specific primers) by polymerase extension or DNA ligation for both global and targeted transcriptome profiling. Like a previous method (*24*), PIXEL-seq can be applied to spatial proteomics and simultaneous multi-omics (e.g., RNA and protein) assays by using DNA-barcoded affinity reagents, such as antibodies and nucleic acid aptamers. Together, continuous polony gels represent a new generation oligo array which achieves one of the highest feature densities and homogenous oligo distributions, providing an ideal platform for spatial molecular barcoding in complex tissues.

## Methods

### Spatial template DNA construction

Spatial template sequence was synthesized by Integrated DNA Technologies. The template was PCR amplified (Taq 2x master mix, NEB M0270; Bridge primers; 15 cycles) and size selected by 2% agar gel.

#### Template sequence

AATGATACGGCGACCACCGAGATCTACACGACGCTCTTCCGATCTNNNNNNNNNNNNNNNNNNNNNNNAGAGAATGAGGAACCCGGGAACAATGATGGAATTTTTTTTTTTTTTTTTTTTTCGATCGACCACCGAGGTTGCCGGACTAGCGCAAGTACTTGTCCATTCCTGAAGAAATATTATATTTATACAACTTACCCATAGAATCCTATTTACTAGGAAAGGAAAAGCCTCCTATTTATACGAAGGTTGTAGAGCTTTCTCAACAACAGTGGAATATCAATGATAGAACAATTGCCGATGTATTAGATGGGGTCTTAATAACACCTCGATTCGAATCGAGATCTCGTATGCCGTCTTCTGCTTG.

Bridge amplification forward primer (BA(+)): AATGATACGGCGACCACCGAGAUCTACAC. Bridge amplification reverse primer (BA(-)): CAAGCAGAAGACGGCATACGAGAT.

### PAA gel casting

PAA gel casting was performed similarly as previously reported (*14*). A major modification is to seed templates on to the PAA gel surface instead of embedding them into the gel, so polonies can mostly form on the gel surface. The gels were casted on round coverslips (Warner Instrument 64-1693), which was assembled into a FCS2 flow cell (Bioptechs) for amplification and sequencing.

### Polony amplification

Polony amplification was performed similarly as previously reported (*14*). Typically, 35 amplification cycles were performed before the *Taq*I digestion.

### *Taq*I digestion

To generate single stranded DNA (ssDNA) for polony sequencing as well as exposing the poly(T) probe for reverse transcription, polony gels were digested with 800 unit/mL (final concentration) *Taq*I (NEB R0149) diluted in 1× CutSmart buffer at 60°C for 1 hour.

### Polony Sequencing

Single-stranded polonies were hybridized with a 2 μM sequencing primer (CTGCGGCCGCTAATACGACTCACTATAGGGATC) in a 5×SSC hybridization buffer. The sequencing was performed on an in-house constructed automated sequencing platform at 60°C using Illumina HiSeq SBS Kit v4.

### Image acquisition

Images were acquired using a Nikon Ti-E automated inverted fluorescence microscope equipped with a Perfect Focus System (PFS), a Nikon CFI60 Plan Fluor 40×/1.3-NA Oil Immersion Objective Len, a linear encoded motorized stage (Nikon Ti-S-ER), and an Andor iXon Ultra 888 EMCCD camera (16-bit dynamic range, 1,024×1,024 array with 13 µm pixels). Two lasers (Laser Quantum GEM, 532 nm, 500 mW and Melles Griot 85-RCA-400, 660nm 400 mW) were used for four-color sequencing imaging.

### Gel priming with poly(T) or gene specific capture probe(s)

For a global transcriptome analysis, we directly used the template with a poly(T) sequence adjacent to a *Taq*I restriction site. To add designed probe sequence(s) to polony gels for capturing specific transcripts, we synthesized 58-bp oligos bearing 8 random bases as unique molecular identifier (UMI) and a probe sequence. The oligos (10 μM) were annealed to restriction digested gel oligos bearing a universal adaptor in the hybridization buffer at 85°C for 6 min followed by decreasing the temperature to 40°C and washing off unbound primers with amplification buffer. The extension was performed with the Bst polymerase at 60.5°C for 5 min. After extension, polony gels were washed with 2 mL elution buffer (1× SSC, 70% formamide) to generate single-stranded capture probes. Primed polony gels are stored in 100% formamide.

### Library construction from tissue sections

A sequenced polony gel was pre-coated with nucleus staining buffer (0.1×SSC, 2.5× SYTOX Green or 2.5× TO-PRO-3 iodide, 0.4× RT buffer). Frozen tissue was placed at -20°C in a Cryostat NX70 (Thermo Scientific) for 15 minutes. Tissue was then mounted onto the cutting block with OCT and sliced at 10 µm thickness. Tissue sections were then transferred on the surface of the polony gel attached on a coverslip. The coverslip was then reassembled back to a FCS2 flow cell. The tissue hybridization buffer (6×SSC, 2U/µL RNAseOUT) was gently injected into the flow cell, and the flow cell was incubated for 15 min at room temperature for RNA hybridization. First-stand cDNA synthesis was performed in the RT solution for 15 minutes at 37°C and then 45 minutes at 42°C. A RT reaction comprises 29 µL H_2_O; 13 µL Maxima 5× RT buffer (Thermofisher, EP0751); 13 µL 20% Ficoll PM-400 (Sigma, F4375); 6.5 µL 10 mM dNTPs; 2 µL RNaseOUT (Thermofisher, 10777019); 1.75 µL 50 µM template switch oligo (Qiagen 339414YCO0076714); 3.25 µL Maxima H-RTase (Thermofisher, EP0751)).

### Tissue cleanup

72 µl Proteinase K digestion solution (63 uL Qiagen PKD buffer, 9 uL Proteinase K) was injected into the flow cell and incubated at 55°C for 30 min. 2 mL of the elution buffer was applied to remove genomic DNA and other contaminants from the polony gel.

### cDNA amplification

To recover spatially indexed cDNA in a polony gel, we performed PCR amplification by transferring the polony gel into a custom-made chamber on a slide and inserting it to a DNA Engine Slide ChambersAlpha unit (Bio-Rad) for amplification. After 2-cycle PCR, the products were purified by Ampure XP beads (Beckman Coulter) with 0.6× beads-to-sample ratio. The purified products were further amplified to obtain 5-10 ng DNA per sample. Typically, we performed 11 PCR cycles for a 4 mm^2^ tissue section. A cDNA amplification mix comprises 14 µL H_2_O, 25 µL 2× Q5 master mix (NEB); 1 µL 10 µM TruSeq PCR handle primer (IDT), 1 µL 10 µM TSO PCR primer (IDT) and 7 uL perified DNA.

### Library construction

DNA concentration was measured by Qubit 4 (Thermo Fisher). The library was constructed using an Illumina Nextera kit following the manufacture’s protocol.

### Estimation of diffusion

Pixel-level intensity matrices were extracted from a nucleus-stained image and a Cy5-dCTP labelled cDNA image captured from the same tissue section. Ten small regions were calculated to determine the transformation matrix and the scaling factor using Matlab build-in function, imregister().

### Pixel-level base calling

After the sequencing images per channel per cycle were collected, we extracted pixel-level intensities and divided into 16 groups for each one. The intensity matrix was loaded into 3Dec (*25*) for base calling separately. We then bin the dataset to obtain one unique spatial index list based on the following rule: (1) the neighbor pixel (< 3 pixel) should have more than two mismatches, otherwise they will be merged into one. (2) calculate the correlation coefficient (cor) between the sequence-unassigned pixel and its neighbor pixel, assign the same spatial index if cor is more than 0.8.

### Bead-shaped spatial indexed transcriptome simulation

In order to better understand the impact of feature size on assigning mapped reads to unique cells, we used the recent published seqFISH+ data1 (pixel size = 0.103µm) to simulate a spatial bead-indexed transcriptome experiment. The simulation used bead-shaped indexing features with 5% variation in radius through Gaussian distribution. On a 2D plane, a series of simulated beads with different diameters (10 to 1 µm) are progressively packed and linked to transcripts within its radius based on their coordinates. The simulation was run 10 times using a raw dataset (average cell diameter is 15.9 µm) and a modified dataset (the average cell diameter was decreased to 7.95 µm).

### PIXEL-seq data analysis

We first extracted the spatial index sequences and UMIs from the read one FastQ files by Cutadapt. We then used bowtie to map the index sequences back to predetermined polony spatial indices with less than one mismatch. The corresponding read two files were aligned to mouse transcriptome (GRCm38) using STAR with default setting. The sequencing reads with the same mapping locus, UMI, and spatial index will be collapsed into one record.

### HiPlex FISH

Mice were anesthetized with Beuthansia (0.2 ml, i.p.; Merck) then brains were rapidly removed and flash frozen on dry ice. Coronal sections (10 μm) were cut on a cryostat (Leica Microsystems) and mounted on Superfrost Plus slides (Fisher). RNAScope HiPlex assay (ACDBio) was performed following the manufacturer protocol. Briefly, slides were fixed in 4% paraformaldehyde, dehydrated in 50%, 70%, and 100% ethanol, then treated with Protease IV. Probes (Penk, Pdyn, Calca, Tac1) were hybridized at 40°C for 2 h then amplified for detection. Cell nuclei were stained with Dapi then fluorescent images were acquired using a Keyence BZ-X700 microscope. Images were registered using HiPlex image registration software (ACDBio) and minimally processed in Fiji (ImageJ) to enhance brightness and contrast for optimal representation of the data.

## Acknowledgements

We thank B. Wang and W. Liang for contributions to polony sequencing data analysis and D. Sun and J. Cheng for setting up the sequencing platform. This work was supported by a startup fund of the University of Washington (to L.G.).

