## Supplemental Figures and Table for "Continuous Polony Gels for Tissue Mapping with High Resolution and RNA Capture Efficiency"

**Supplementary Figure 1.**

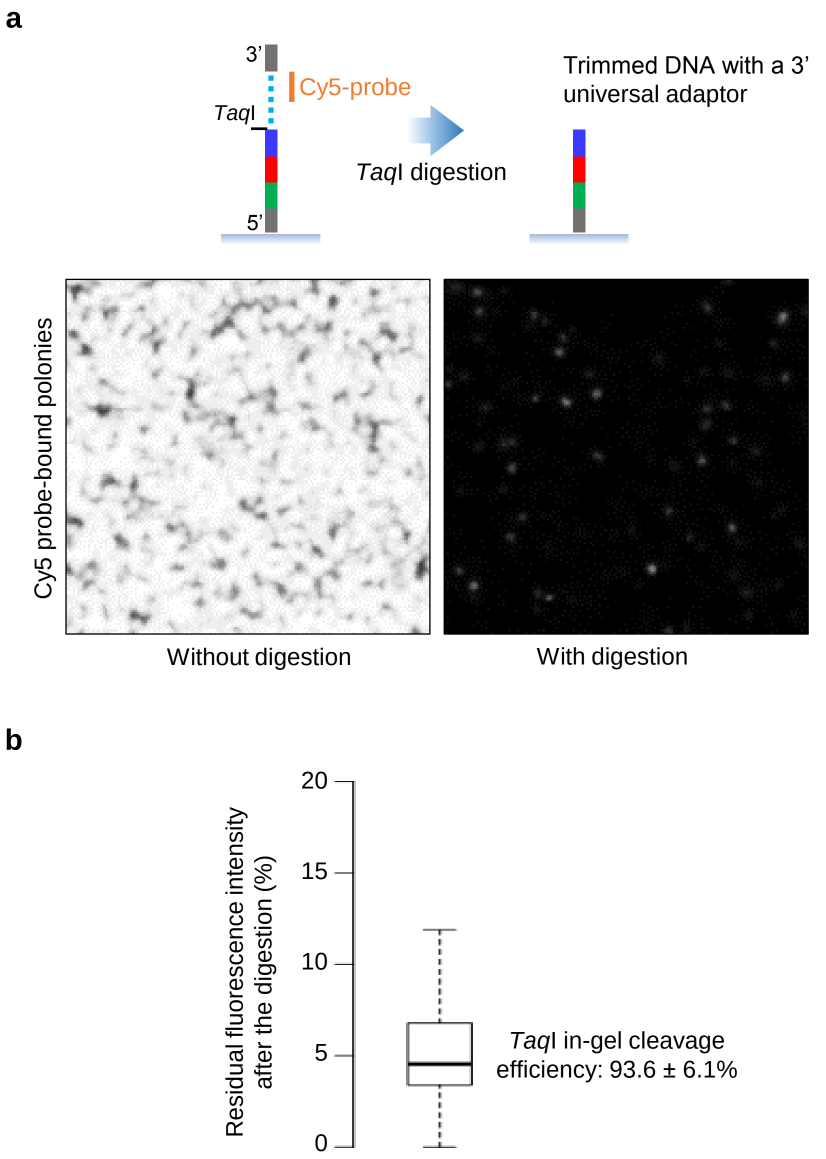


**Figure S1. Analysis of *Taq*I cleavage efficiency of continuous polony DNAs. (a)** Detection of non-cleaved polonies by a Cy5-probe. The digestion was performed at 60˚C for 1 hour and the residual DNAs were hybridized with a Cy5-probe to detect non-cleaved DNAs. **(b)** Quantification of the ratio of cleaved DNA by image analysis. Eight randomly selected fields-of-view (1,024×1,024) for each condition were subjected to image quantification.

**Supplementary Figure 2.**

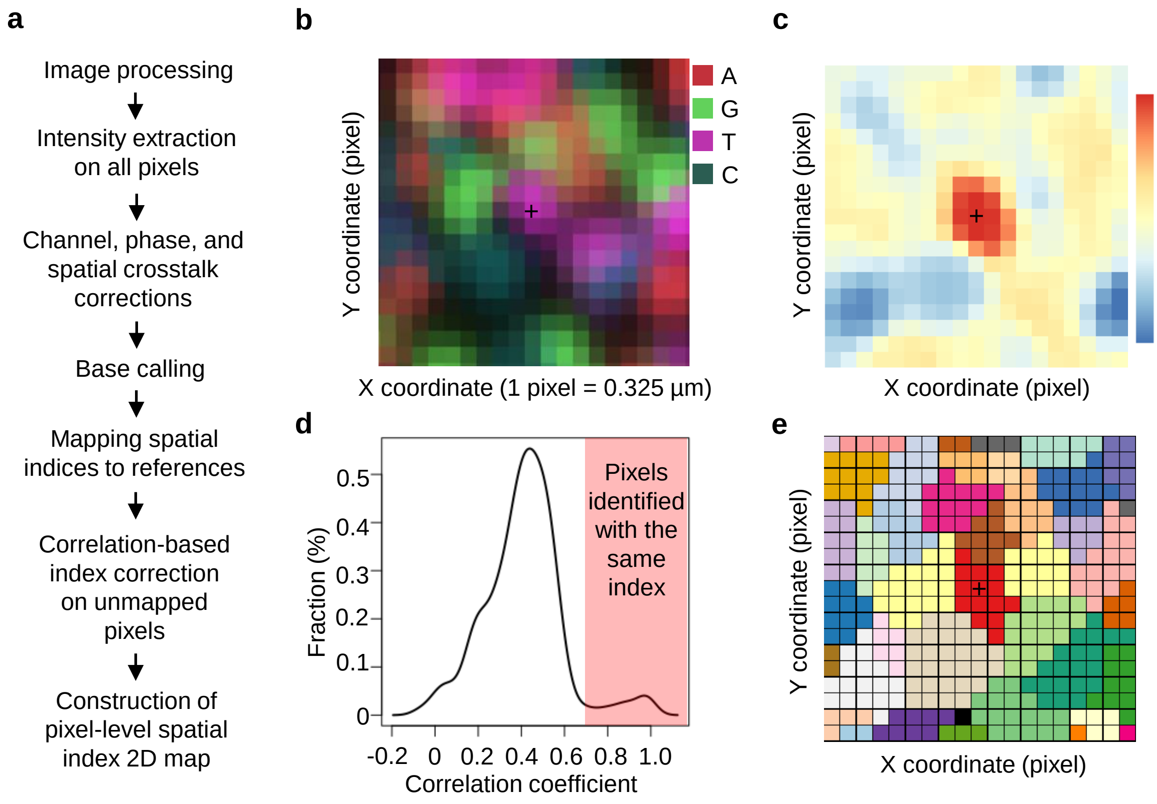


**Figure S2. Pixel-level reconstruction of spatial indices. (a)** Bioinformatic pipeline of pixel-level spatial index map reconstruction. Briefly, the pixel-level raw intensities were extracted from sequencing images for base calling. By analyzing Illumina sequencing reads, we check spiked-in 6-mer reference sequences within the barcodes and discard the unmatched. The remain reads are allocated to their coordinates. Unassigned pixels were directed to the positions by neighbor pixel-intensity correlation (dist = 5 pixels). **(b)** Example of raw image of four-color sequencing. **(c)** Spatial heatmap of correlation-coefficient between a center pixel and its neighbors. **(d)** The density plot of correlation-coefficient. Cutoff = 0.7 was chosen based on the distribution. **(e)** A final reconstructed pixel-level spatial index map.

**Supplementary Figure 3.**

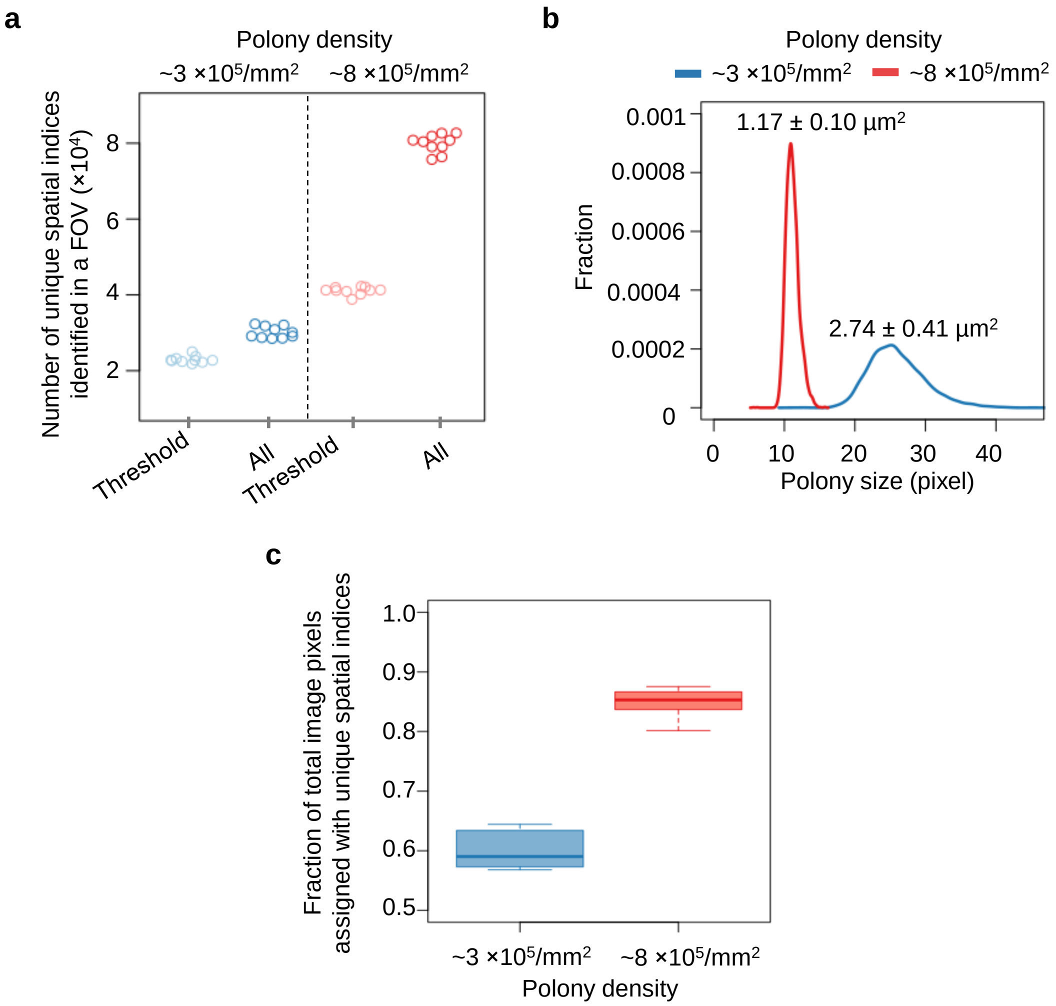


**Figure S3. Pixel-level base calling is required for high-density polonies. (a)** Comparison of threshold-based polony base calling and pixel-level base calling. 10 FOVs were included for the analysis. (**b**) Distribution of estimated polony diameter. $pi \times 1/4d^2 = n \times pixel \times pixel$. (**c**) Comparison of the ratios of confidently called pixels (n = 10).

**Supplementary Figure 4.**

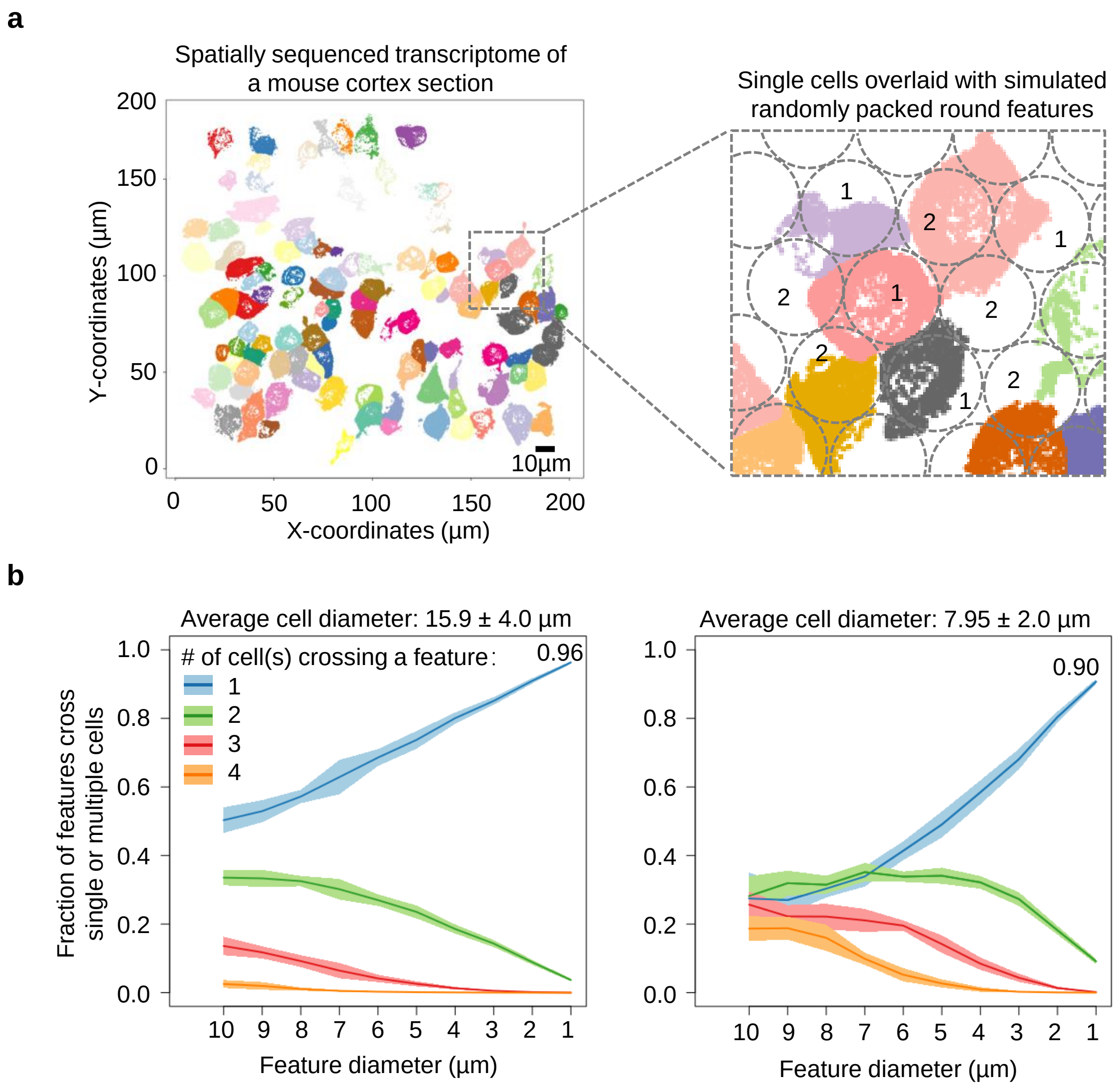


**Figure S4, Simulation of the impact of array feature sizes on single cell isolation efficiency. (a)** A published seqFISH+ data[1] (pixel size = 0.103 µm) was used to simulate a spatially indexed transcriptome experiment. The simulation adopts bead-shaped spatial index with 5% variation in radius through Gaussian distribution. On a 2D plane, a series of simulated beads with different diameters (10 - 1 µm) are progressively packed and linked to transcripts within its radius based on their coordinates. The simulation was run 10 times using the raw dataset (average cell diameter:15.9 µm) and the modified dataset (the average cell diameter was proportional decreased to 7.95 µm). (**b**) Fraction of features that can be confidently assigned to single cell. Cutoff = 0.7, 70% of the mRNAs from single feature belongs to the same cell.

**Supplementary Figure 5.**

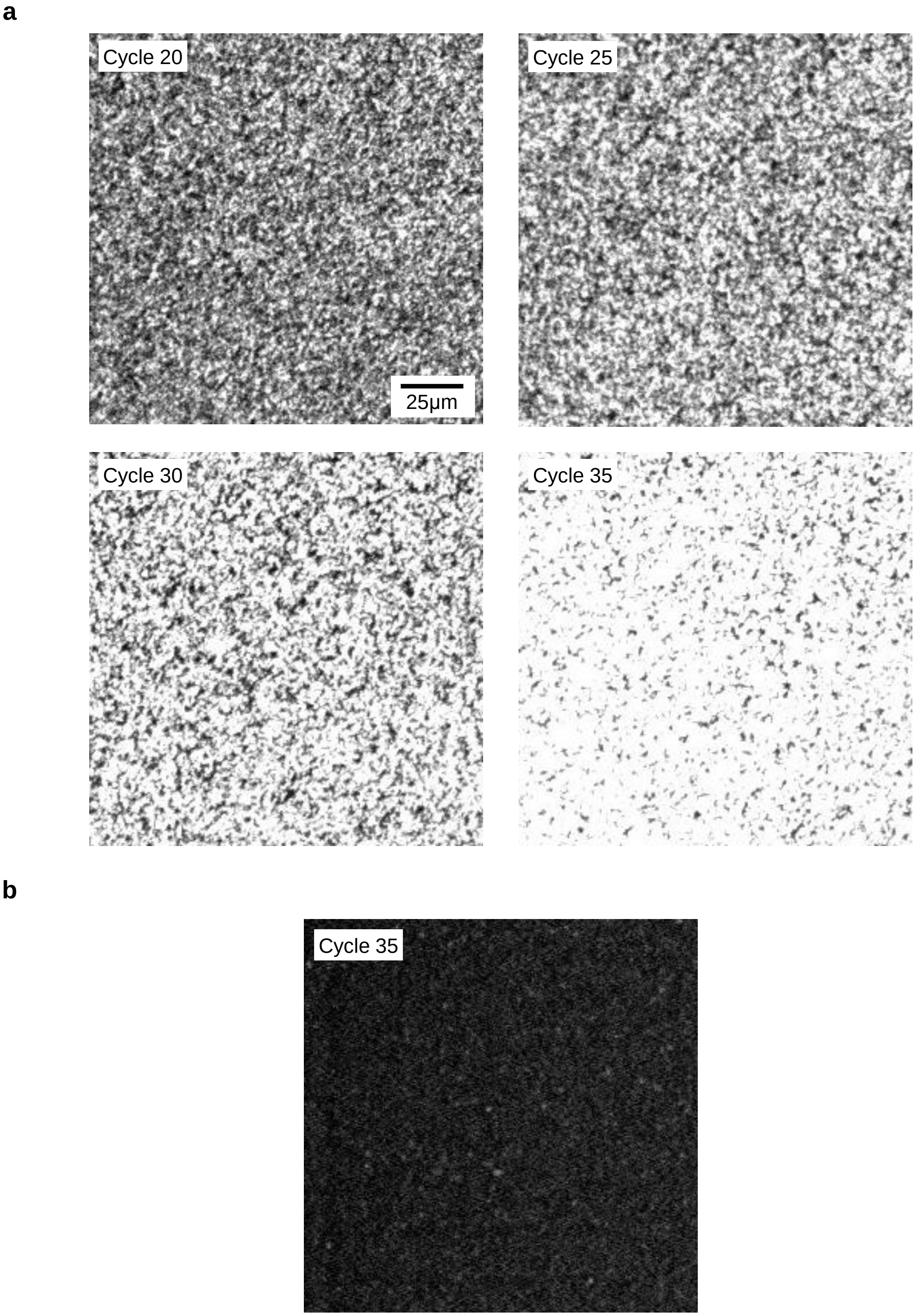


**Figure S5. Real-time analysis of polony intensities in crosslinked (a) and linear PAA (b) gels.** Polonies were stained by SYBR.

**Supplementary Figure 6.**

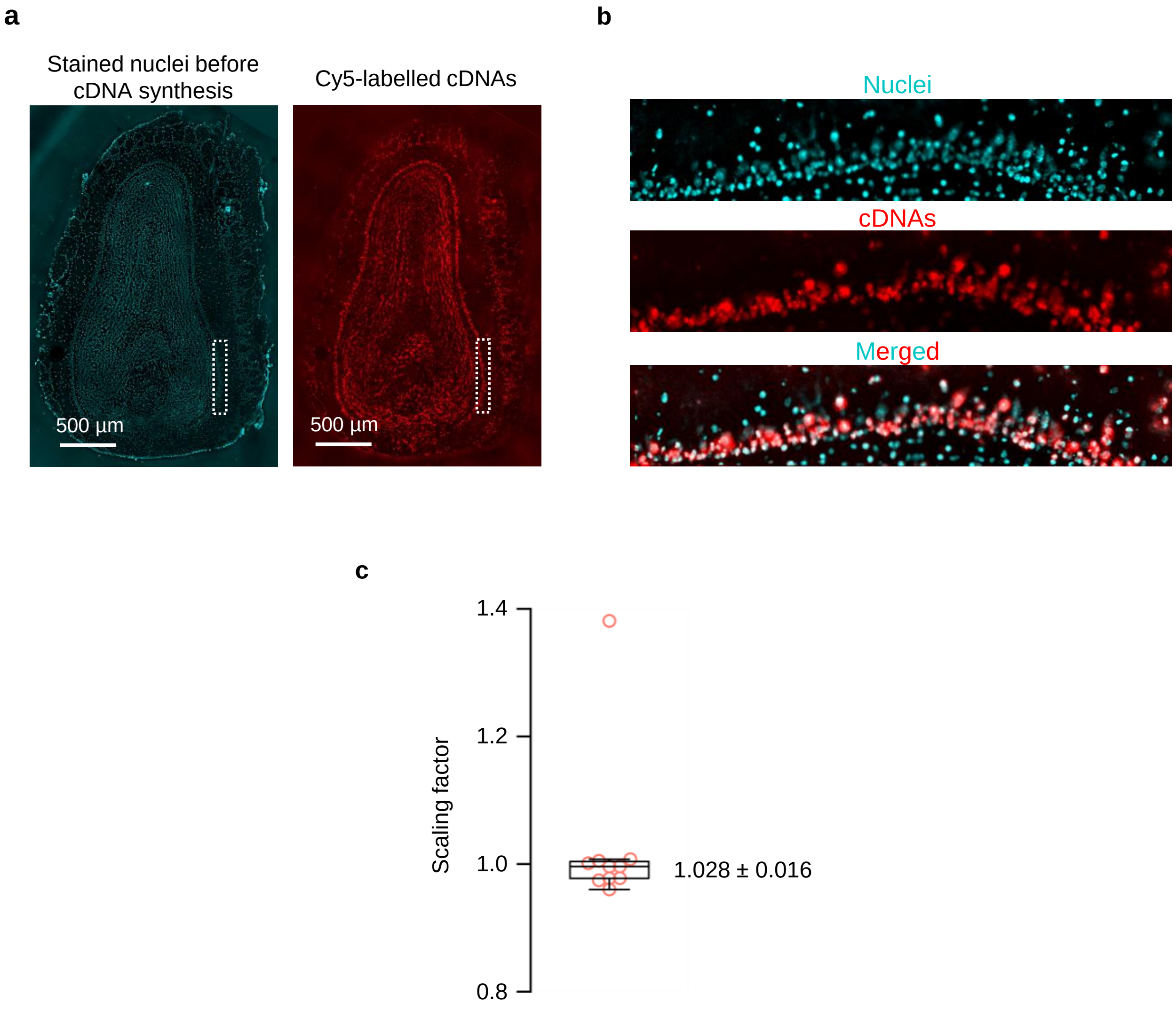


**Figure S6. Analysis of template diffusion in polony gels.** (**a**) Pre-scanned nucleus image of the mouse main olfactory bulb section (SYTOX) and a Cy5-dCTP incorporated cDNA image of the same tissue. (b) Zoom-in visualization of the nuclei, cDNAs, overlaid signals after image registration. (c) Measure of cDNA lateral diffusion in a randomly selected region using a scaling factor as an indicator.

**Supplementary Figure 7.**

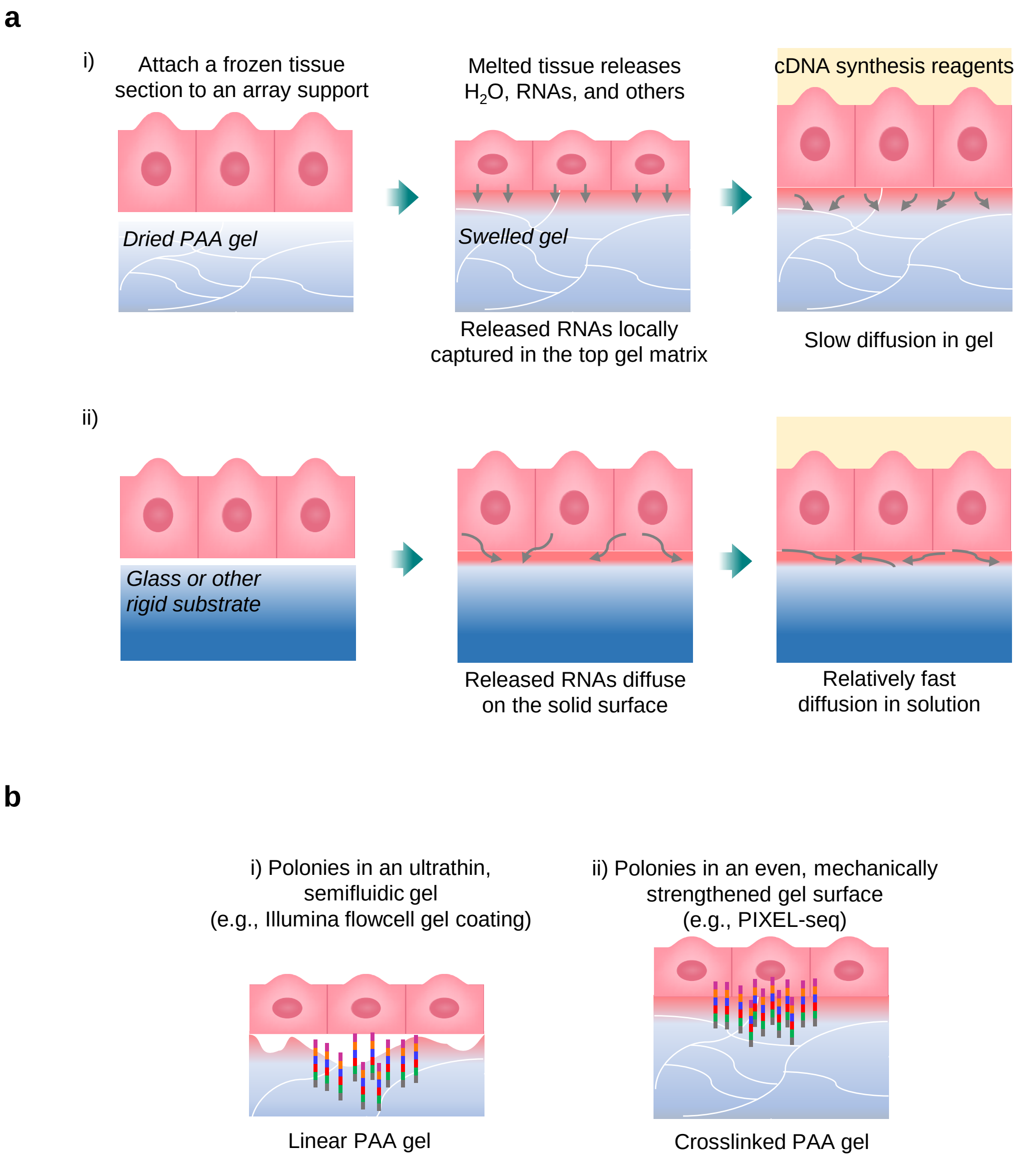


**Figure S7. (a)** Proposed model for polony gel-constrained template diffusion. The gel substrate is compared with solid substrates. (**b**) Proposed model for polony gel-enhanced RNA capture. The crosslinked PAA gel substrate is compared with the linear PAA substrate.

**Supplementary Figure 8.**

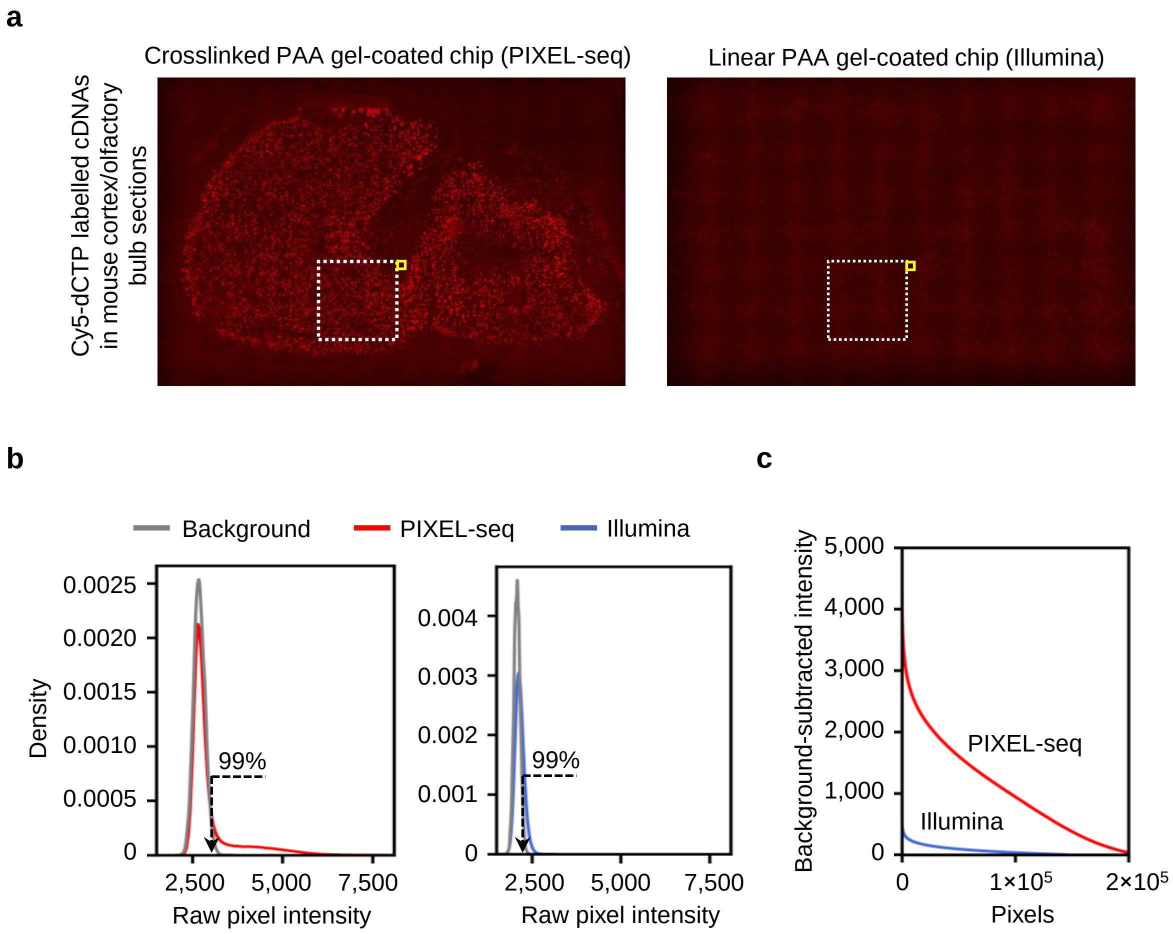


**Figure S8. Comparison of Cy5-labelled cDNAs on crosslinked PAA and Illumina linear PAA gel substrates. (a)** Cy5 images of the mouse accessory olfactory bulb sections on crosslinked and Illumina linear PAA gels. Quantitative comparison of Cy5 signals before (**b**) and after (**c**) the background subtraction.

### Supplementary Figure 9.

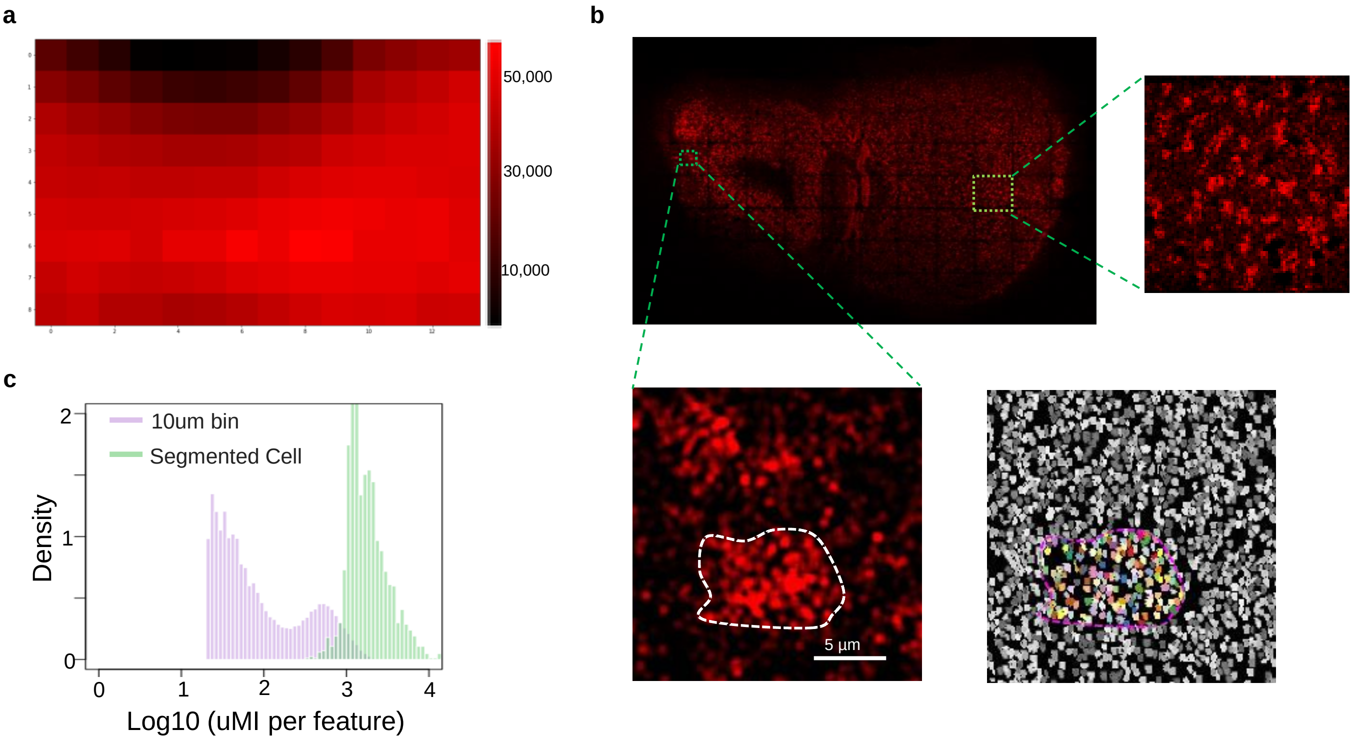


**Figure S9. Spatial gene profiling of the mouse accessory olfactory bulb (aOB) by PIXEL-seq. (a)** Sequenced polony density of the gel used in this assay. **(b)** Spatial distribution of gene density per polony (1 pixel = 0.325×0.325um). Close visualization of subcellular gene abundance distribution (bottom). **(c)** Total UMI distribution per feature. 10-µm bins were generated by grouping 33 pixels. Segmented cells were manually curated using the nucleus-stained image as a reference.

**Supplementary Figure 10.**

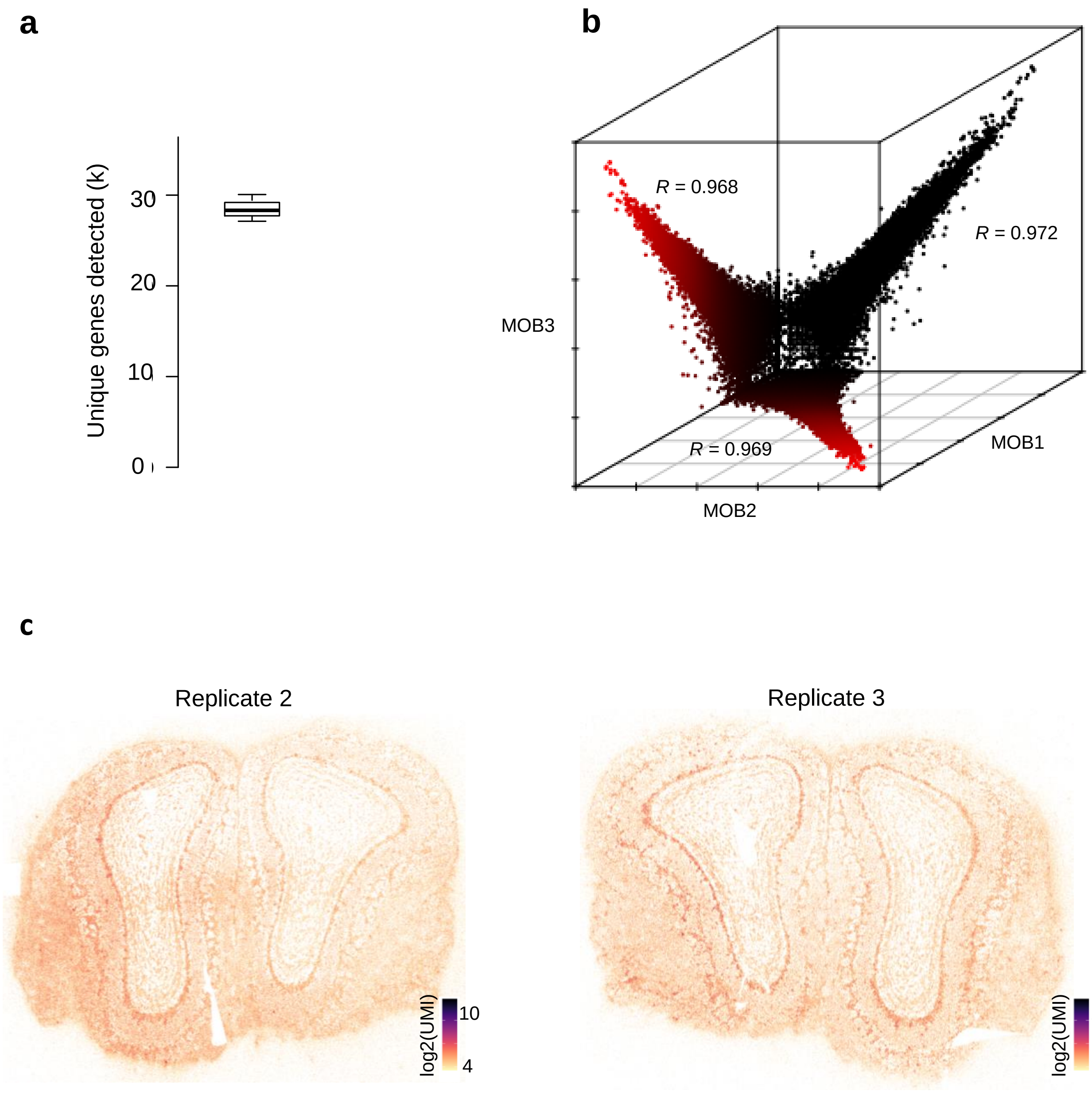


**Figure S10. PIXEL-seq performance on MOB replicates.** (**a**) Boxplot of the total unique genes found in the three MOB replicates. (**b**) Correlation of gene expression profiles for each MOB replicate (n = 3). Denoted are the Pearson correlation coefficients between groups. **(c)** Spatial heatmap of total UMIs per spatial barcode for two MOB replicates.

**Supplementary Figure 11.**

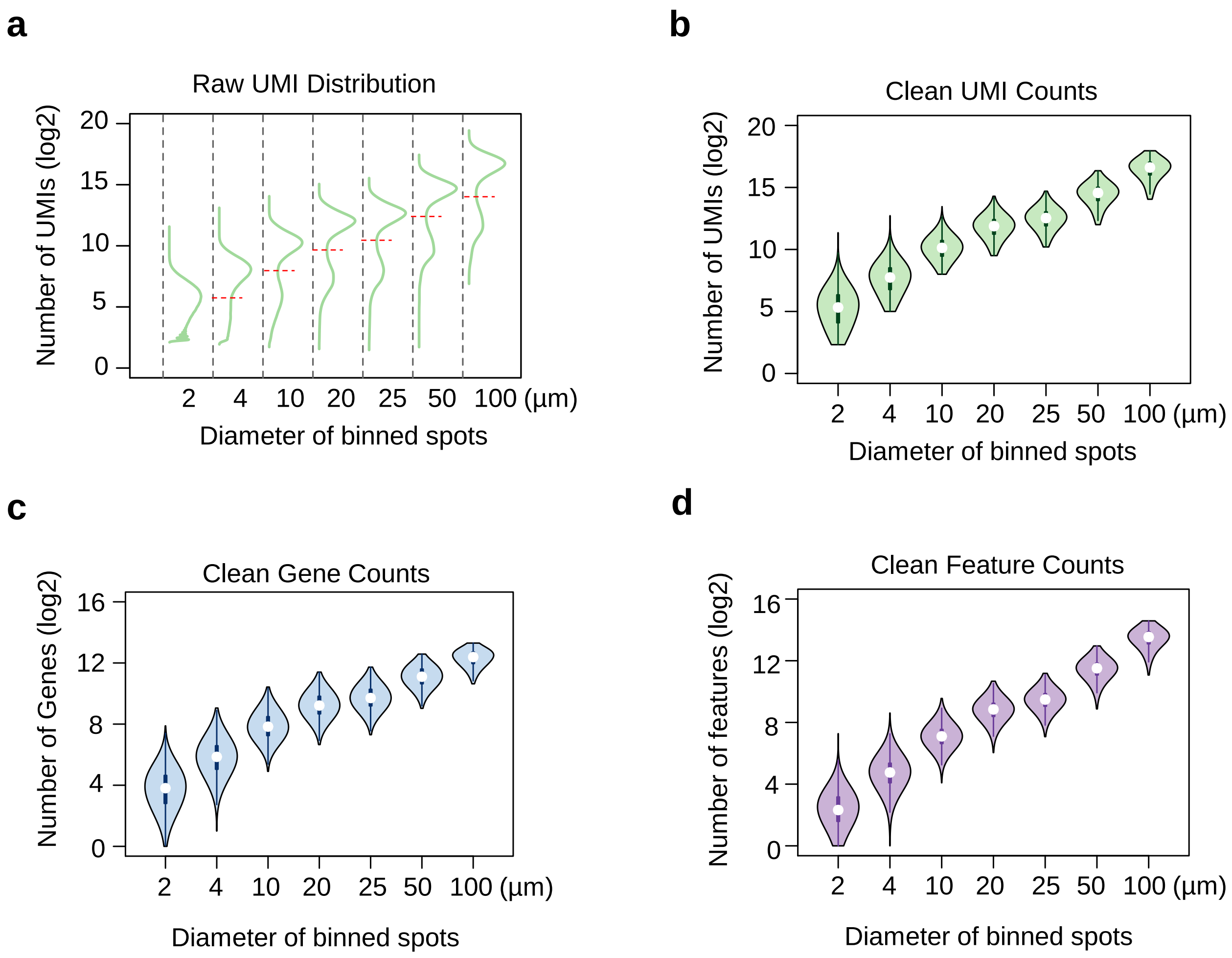


**Figure S11. Feature statistics of mouse main olfactory bulb (mOB) by PIXEL-Seq. (a)** Raw UMI distribution at various binned sizes. Red lines label the cutoffs for each group. The **v**ioplots of clean UMI counts (**b**), clean gene counts (**c**), and clean feature counts at indicated feature sizes.

**Supplementary Table S1.**

| | Spatial resolution | | Detection sensitivity | | |
|---|---|---|---|---|---|
| | Feature size & center-to-center distance (μm) | Feature density (clusters/$mm^2$) | Oligo density (molecules/ $\mu m^2$) | Array area covered by oligos (%) | Array substrate |
| **Visium** | 55 & 100 | $\sim 10^2$ | $\sim 8\times 10^3$ | ~24 | Solid surface |
| **Slide-seq** | 10 & 10 | $\sim 10^4$ | unknown | ~78 | Solid surface |
| **HDST** | 2 & 3 | $\sim 10^5$ | unknown | ~34 | Solid surface |
| **PIXEL-seq** | 1 & 1 | $\sim 10^6$ | $\sim 2\times 10^4$ | >90 | PAA gel |
